# A 13-subunit c-ring in the *Chlamydomonas* chloroplast ATP synthase lowers the H⁺/ATP cost of carbon fixation

**DOI:** 10.64898/2026.09.03.749062

**Authors:** Katarzyna Lorencik, Sebastian Pintscher, Katherine H. Richardson, Hannah Takahashi, Matthew S. Proctor, Michał Rawski, C. Neil Hunter, Andrew Hitchcock, James N. Blaza, Matthew P. Johnson

**Author notes:** These authors contributed equally (Katarzyna Lorencik, Sebastian Pintscher).

## Abstract

The chloroplast F_1_F_o_ ATP synthase is a rotary motor that converts the light-driven proton-motive force into the chemical energy of ATP. The number of c-subunits in its rotor fixes the number of protons translocated per ATP formed, a fundamental parameter of bioenergetic systems. The reference spinach enzyme possesses fourteen c-subunits and a H^+^/ATP ratio of 4.67. Green algae additionally operate a carbon-concentrating mechanism that sustains CO_2_ fixation in water at a substantial cost in ATP, yet the structure of the algal motor, and whether its bioenergetic parameters differ from those of vascular plants, remains unresolved. Here, a 2.2 Å structure of the ATP synthase of *Chlamydomonas reinhardtii* reveals that the enzyme carries a thirteen-membered c-ring, the first departure from c_14_ in a chloroplast, and with a lower predicted H^+^/ATP ratio of 4.33. Ordered waters trace a Grotthuss proton relay through the membrane, where an insulating triad separates the proton loading and unloading sites and couples flux to rotation. A single substitution in the redox switching γ-subunit abolishes the contact with the catalytic β-subunit that idles the enzyme in darkness in vascular plants. These unique features of the algal ATP synthase lower the H^+^/ATP cost of carbon fixation in the light and facilitate acetate metabolism in the dark.

## Introduction

The F_1_F_o_ ATP synthase is a rotary molecular motor that couples the transmembrane proton-motive force (pmf) to the synthesis of ATP, and is found in the energy-transducing membranes of bacteria, mitochondria and chloroplasts^1,2^. The enzyme is built from two linked rotary components. In the membrane-embedded F_o_ domain, protons pass through two offset half-channels in the a-subunit and drive the rotation of a ring of c-subunits, each of which carries an essential carboxylate that is protonated on one side of the membrane and deprotonated on the other^3^. This rotation is transmitted through the central γε stalk to the F_1_ head, where the alternating conformations of three catalytic β-subunits and structurally supporting α-subunits synthesise ATP by the binding-change mechanism^4,5^. A peripheral stalk braces the α_3_β_3_ head against the torque of the rotor.

Because a single revolution of the c-ring yields three molecules of ATP, the number of c-subunits fixes the number of protons translocated per ATP and therefore sets the H^+^/ATP ratio, one of the fundamental bioenergetic parameters of the cell^6^. Ring stoichiometry is not conserved across life, ranging from eight in bovine mitochondria, giving an H^+^/ATP ratio of 2.7, to as many as fifteen in some cyanobacteria, taking the ratio to 5^7–9^. The value adopted by a given organism is thought to match the proton cost of ATP synthesis to the pmf that its membrane can sustain^9,10^. Within the chloroplast lineage, however, high-resolution structural knowledge has until recently been confined to a single vascular plant: the spinach enzyme possesses a 14-membered c-ring and an H^+^/ATP ratio of 4.67, and has provided the reference structure for the chloroplast motor^11,12^.

The H^+^/ATP ratio is of particular consequence in the green algae. Unlike vascular C_3_ plants, *Chlamydomonas reinhardtii* operates a carbon-concentrating mechanism that raises the concentration of CO_2_ around Rubisco and suppresses photorespiration, but at a substantial additional cost in ATP^13^. Since linear electron transfer (LET) alone cannot meet the elevated ATP/NADPH ratio required, the alga generates additional pmf through cyclic (CET), pseudocyclic (PCET) and chloroplast-to-mitochondrion electron-transfer (CMET) pathways that are believed to augment ATP levels^14,15^. The H^+^/ATP ratio of the chloroplast ATP synthase sits at the centre of this energy balance, because it determines how much pmf must be spent for each molecule of ATP. Recently, a structure of the *C. reinhardtii* ATP synthase determined using cryogenic electron microscopy (cryo-EM) has been reported, though the low spatial resolution, particularly in the F_o_ domain, leaves these crucial questions unanswered ^16^.

*C. reinhardtii* also regulates the chloroplast ATP synthase differently in the dark^17^. In vascular plants a redox switch on the γ-subunit forms a disulphide in darkness that raises the proton-motive-force threshold for activity and idles the enzyme to conserve ATP^11,12^, whereas in *C. reinhardtii* this thiol modulation is functionally disconnected, keeping the ATP synthase active in the dark to support metabolic coupling between the chloroplast and its acetate-fuelled mitochondria^17–19^. Currently, there is no structural basis for these differences.

Here, we sought answers to these outstanding questions by using cryo-EM to determine the high-resolution structure of the chloroplast ATP synthase of *C. reinhardtii*.

## Results

### Overall architecture and a 13-membered c-ring

We treated *C. reinhardtii* thylakoid membranes with the detergent α-dodecyl maltoside, separated the solubilised complexes by sucrose gradient ultracentrifugation, and determined the structure of ATP synthase by single-particle cryo-EM (Fig. 1a; Extended Data Fig. 1). A consensus reconstruction at 2.2 Å, supplemented by focused refinements of the F_1_ head, the central stalk, and the F_o_ membrane region, allowed the complete enzyme to be built (Extended Data Fig. 2; Extended Data Table 1). The structure contains the expected chloroplast subunit complement: an α_3_β_3_ catalytic head, a central γε stalk, a peripheral stalk formed by the b and b’ subunits together with δ, and a ring comprised of the c-subunits and the a-subunit (Fig. 1a-d).

**Fig. 1.**
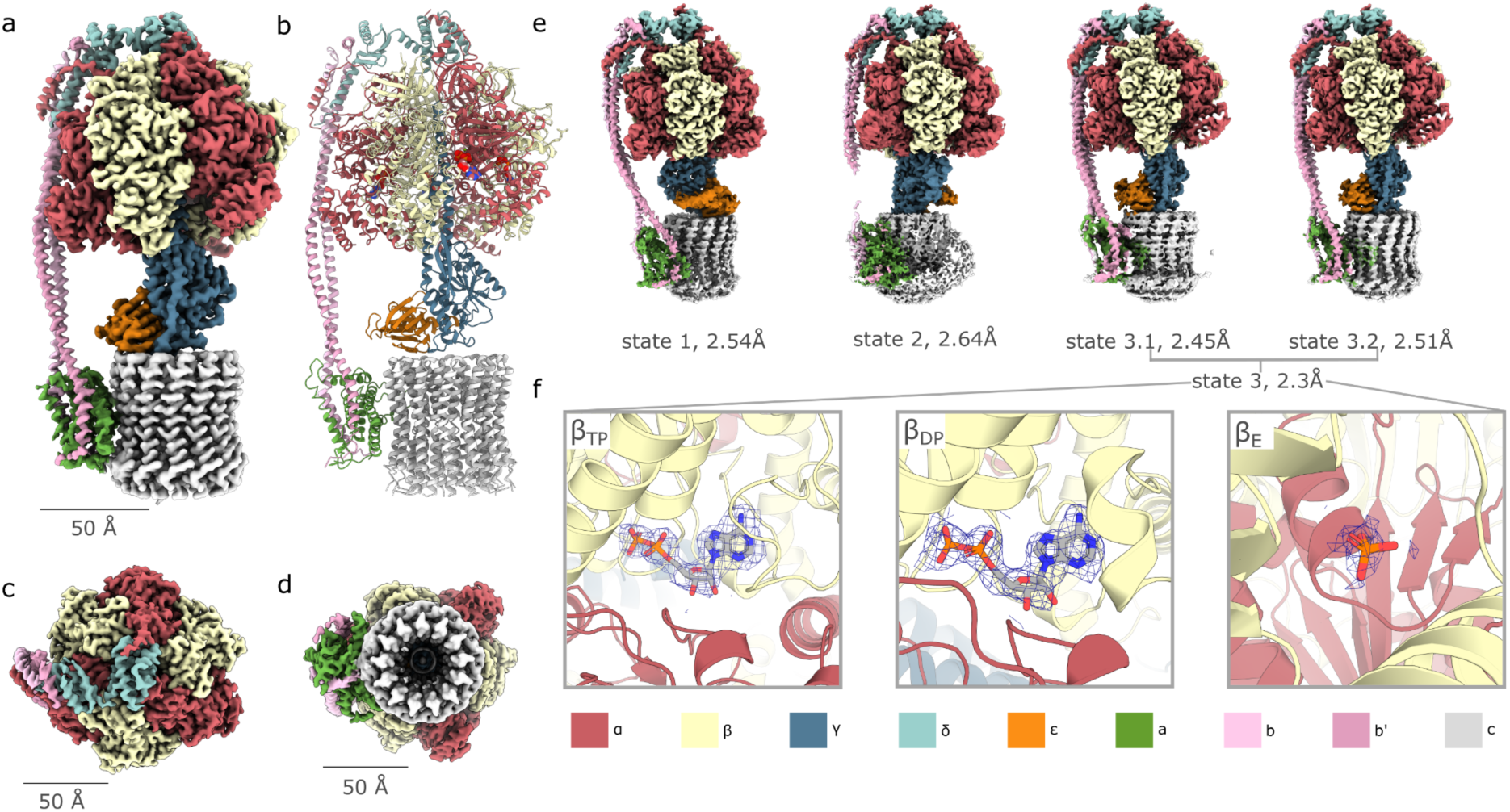
Architecture and rotary states of the *C. reinhardtii* chloroplast ATP synthase. ***a***, Composite cryo-EM density of the intact enzyme at 2.2 Å with subunits labelled (α_3_β_3_, γ, ε, δ, a, b, b’ and the c_13_ ring). ***b***, Modelled subunits of ATP synthase. Composite cryo-EM density view from ***c***, the top of the F1 domain and ***d***, the bottom of the F_o_ domain. ***e***, Cryo-EM maps of the three principal rotary states (dwells 1–3) of the catalytic head, related by successive ∼120° rotations of the γ-subunit, together with the two substates of state 3. ***f***, Nucleotide/ phosphate occupancy of the three catalytic sites (β_TP_ = tight, β_DP_ = loose, β_E_ = empty) defined by the binding change mechanism within the F_1_ domain.

Saliently, the c-ring of the *C. reinhardtii* enzyme is built from thirteen c-subunits, rather than the fourteen found in spinach chloroplast ATP synthase (Fig. 1d). The thirteen protomers are related by a mean rotation of 27.7° and form a closed, symmetric ring. A 13-membered ring translocates thirteen protons per revolution and therefore gives a predicted H^+^/ATP ratio of 4.33, lower than the 4.67 of the spinach c_14_ enzyme, and establishes that the proton cost of ATP synthesis differs between the algal and vascular-plant chloroplasts.

### The three catalytic dwells of the rotary cycle

The catalytic head is built from three αβ pairs arranged alternately around the central γ-subunit, with the three catalytic nucleotide-binding sites lying at the α/β interfaces and three non-catalytic sites at the complementary interfaces; rotation of the γε rotor drives each catalytic site in turn through the open, loose and tight conformations of the binding-change mechanism^4,20^. Three-dimensional classification (Extended Data Fig. 3) of the F_1_ head resolved the enzyme in three principal rotary states, together with two closely related substates of one of them (Fig. 1e; Extended Data Fig. 4). When the models are superimposed on the stator, the three principal states are related by successive rotations of the γ-subunit of ∼120° (measured as 122°, 120° and 118° between the pairs), and therefore correspond to the three catalytic dwells of a single revolution of the motor (Extended Data Figs 5 and 6). The two substates differ from one another by a rotation of the ε-subunit by 20° with respect to the γ-subunits, thus likely representing a rotational sub-step within the third dwell rather than a distinct catalytic state.

The nucleotide occupancy of the catalytic sites is conserved across the three dwells and is characteristic of a resting enzyme. Two of the catalytic sites contain ADP and the third contains only phosphate, together with associated Mg^2+^, while the γ-subunit sets the register of the open, loose and tight β-conformations around the ring (Fig. 1f and Extended Data Fig. 5). This configuration, with ADP and phosphate but no bound ATP at the catalytic sites, corresponds to the ADP-inhibited resting state described for the oxidised spinach enzyme^11,12^, and contrasts with the ATP-loaded catalytic sites seen when bacterial ATP synthases are supplemented with nucleotide^21^ or when the reduced spinach ATP synthase binds the inhibitor tentoxin^11^. The three non-catalytic sites on the α-subunits are occupied predominantly by ATP; in each state one of these sites shows weaker density, which we have modelled conservatively as phosphate, and the position of this site advances by one αβ pair between successive states, mirroring the 120° stepping of the rotor (Extended Data Fig. 6).

### A localised steric basis for the reduced stoichiometry

The c_13_ ring closely resembles the spinach c_14_ ring in its overall dimensions. The mean radius of the subunit centres of mass is 21.2 Å in *C. reinhardtii* and 21.6 Å in spinach (PDB 6FKF)^12^, so that the algal ring accommodates one fewer protomer within a circumference that is only marginally smaller (Fig. 2a). Because the two mature c-subunits share approximately 79% sequence identity and adopt indistinguishable helical-hairpin folds, the change in symmetry is not apparent from sequence (Fig. 2b) or fold alone, and we sought its origin by comparing the subunit interfaces directly.

**Fig. 2.**
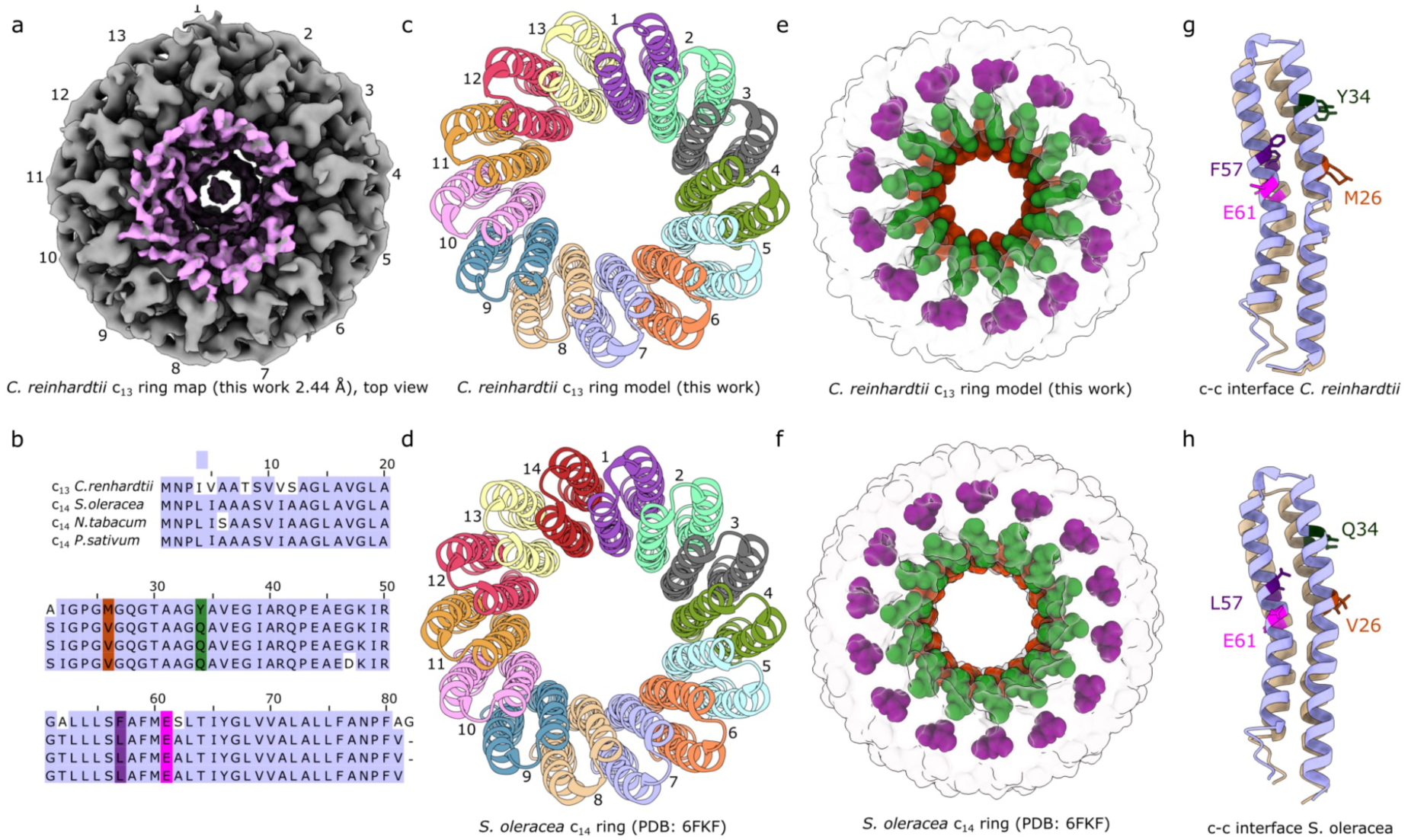
Structural comparison of the *C. reinhardtii* c₁₃ and spinach c₁₄ rings. ***a*,** Top-down view of the *C. reinhardtii* c₁₃ ring from the 2.44 Å c-ring focused refinement, looking along the rotor axis. The associated lipid densities are shown in pink. ***b*.** Alignment of the chloroplast c-subunit sequences from *C. reinhardtii* compared to three vascular plants. The equivalent residues by position are marked with colours and are also highlighted in panels g and h. ***c*, *d*.** c-ring models of *C. reinhardtii* and *S. oleracea* (PDB 6FKF)^12^, respectively, with each of the thirteen c-subunits shown in a distinct colour, and the additional fourteenth subunit of the spinach complex shown in red. The mean subunit centre-of-mass radius differs by 0.44 Å between the two rings (21.19 Å vs. 21.63 Å). ***e*, *f*.** Surface rendered view of the c-ring in e, *C. reinhardtii* and f, *S. oleracea* highlighting the c-subunit side chains of the three non-conserved interfacial residues as space filling models and coloured by position: Met26/Val26 (orange), Tyr34/Gln34 (green), and Phe57/Leu57 (purple). The tighter packing due to the bulkier residues Met/Tyr/Phe in *C. reinhardtii* do not permit addition of a fourteenth subunit in the c-ring. ***g*, *h*.** A single c–c interface extracted from each ring, rendered in identical orientation following structural superposition of residues 30–75 on the c-subunit of *C. reinhardtii* to that of *S. oleracea*. The two subunits of the pair are coloured pale blue and wheat. The relative position of the conserved Glu61 residue is also shown in pink.

The decrease from fourteen to thirteen subunits is accompanied by a widening of the packing between neighbours. The mean centre-to-centre spacing of adjacent protomers is 0.52 Å greater in *C. reinhardtii* than in spinach (10.14 versus 9.63 Å), which widens the angular step from 25.7° to 27.7°; at this spacing a fourteenth subunit cannot be closed into the ring at a comparable radius (Fig. 2c-f). Three non-conserved positions at the subunit interface account for the additional spacing. At positions 26, 34 and 57 the *C. reinhardtii* c-subunit presents methionine, tyrosine and phenylalanine in place of the valine, glutamine and leucine found at the equivalent positions in the spinach protein; in each case the algal residue projects a bulkier side chain into the interface between neighbouring protomers (Fig. 2e-h). Indeed, the Met26 and Tyr34 residues in *C. reinhardtii* fall adjacent to the conserved glycine repeat Gly23xGly25xGly27xGly29xxxGly33, previously identified as crucial to c-subunit packing and thus ring size in ATP synthases^22^. The essential proton-binding glutamate, Glu61, is also conserved in *C. reinhardtii* and occupies the equivalent inward-facing orientation observed in spinach. Consistent with the wider packing, we find the buried surface area at each c–c subunit interface is approximately 3% smaller in *C. reinhardtii* than in spinach (3,498 Å^2^ versus 3,608 Å^2^) (Extended Data Table 2).

The lower stoichiometry of the algal c-ring therefore arises not from a global change in the fold or dimensions of the c-subunit, but from the cumulative steric effect of three larger side chains at the subunit interface. A change at the level of single residues widens the ring by half an angstrom per subunit and lowers the H^+^/ATP ratio of the enzyme. The near-threefold symmetry of the F_1_ head and the 13-fold symmetry of the c-ring are thus mismatched, so that a non-integer number of c-subunits, and hence protons, passes the a-subunit during each catalytic step, on average 4.33 for the c_13_ enzyme. The steps between the three resolved states (Fig. 1e) are nonetheless close to equal, passing 4.4, 4.3 and 4.3 c-subunits, in contrast to the markedly uneven spinach c_14_ enzyme steps (103°, 112° and 145°; 4.0, 4.4 and 5.6 c-subunits)^12^. Following the thermodynamic treatment of Hahn *et al.*^12^, and taking the free energy of ATP synthesis under chloroplast conditions as approximately 51 kJ mol^-1^, each translocated proton contributes about 11.8 kJ mol^-1^, so that the c_13_ steps carry near-uniform free-energy increments of roughly 52, 51 and 50 kJ mol^-1^, against the 44, 48 and 61 kJ mol^-1^ estimated for spinach. A more even division of the driving energy between steps would lower the step-to-step variation in torque that the elastic central and peripheral stalks must accommodate over each revolution.

### An ordered-water relay in the F_o_ proton pathway

To obtain a detailed view of the proton pathway within the F_o_ domain, we performed a local refinement with a tight mask over the c-ring and a-subunit for particles in state 3 (see Extended Data Fig. 1d). The obtained map resolves thirty-four ordered water molecules within the F_o_ domain, both in the aqueous half-channels of the a-subunit and along the interface between the a-subunit and the c-ring (Fig. 3a). These waters form a continuous, hydrogen-bonded chain that links the lumenal entry channel to the essential Glu61 carboxylate of the c-ring (Fig. 3b and c). Such an arrangement is consistent with proton transfer by a Grotthuss-type relay, in which the proton is passed along a hydrogen-bonded water wire rather than carried by diffusion, mirroring and extending the ordered-water network recently visualised in the bacterial ATP synthase of *Pseudomonas aeruginosa*^21^.

**Fig. 3.**
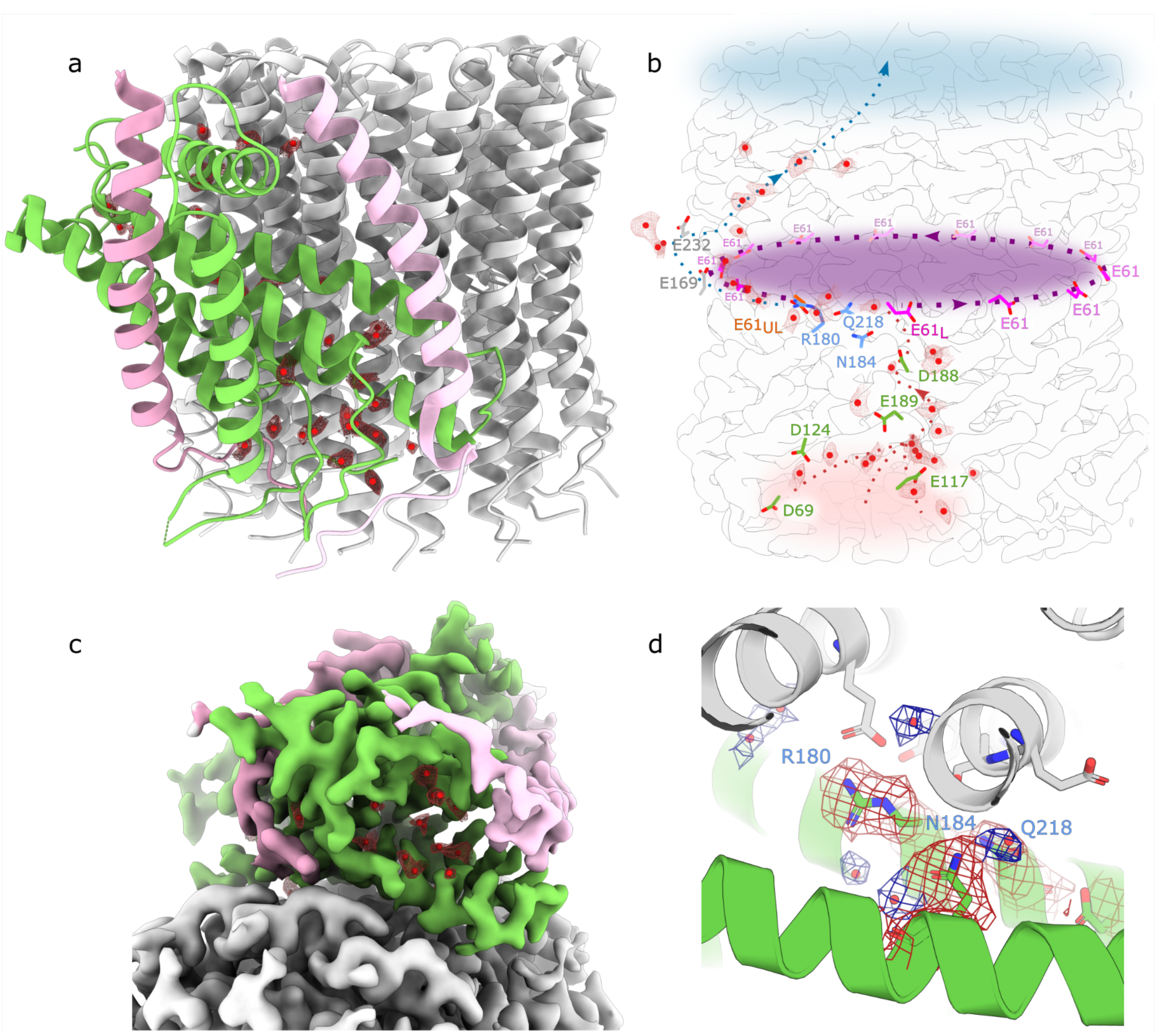
Proton translocation within the *C. reinhardtii* chloroplast F_o_ domain. *a,* F_o_ model shown in a perpendicular view. The water molecules are shown as red dots. ***b,*** Proposed proton transfer pathway within F_o_. Water densities shown as red mesh, c-ring map shown as contours. The protonated Glu61 residue (E61_L_ - Load) is shown in pink, and the deprotonated Glu61 residue (E61_UL_ - Unload) is in orange. The three amino acid residues that prevent a short circuit (the ‘insulating triad’) are shown in blue. Shading denotes the lumenal-facing proton entry vestibule (red), the ring formed from adjacent c-subunit Glu61 residues (purple) and the stromal-facing proton exit region (blue). ***c***, Closeup view of the proton entry half-channel forming a hydrated crevice as viewed from the lumenal side of the membrane. Map coloured according to the subunit colours in Fig 1., with water densities shown as red mesh and water molecules modelled as red dots. ***d*,** Closeup view of the insulating triad. Water densities, and Arg180, Asn184 and Gln218 are shown as mesh.

The conserved Arg180 lies between the two half-channels of the a-subunit, and separates the proton entry and exit pathways (Fig. 3b-d)^12^. Together with Asn184 and Gln218, this residue forms an “insulating triad” that prevents a proton from passing directly between the half-channels without traversing the c-ring, enforcing strict coupling of proton flux to rotation (Fig 3d). The insulating triad allows the loading and unloading sites to be very close together, such that the proton must follow nearly the full 360° of rotation once on the c-ring, ensuring that the maximal torque is extracted from proton flux through the membrane. The positions of these residues are essentially identical in the algal and spinach enzymes, indicating that the proton-relay machinery of the chloroplast motor is structurally conserved even where the c-ring stoichiometry is not.

### The γ-subunit redox loop retains the oxidised conformation but lacks the DELSEED chock in Chlamydomonas

The ∼40-residue ‘redox-loop’ insertion that distinguishes the chloroplast γ-subunit from its mitochondrial counterparts folds into two β-hairpins that project from the central shaft toward the conserved DELSEED motif within the β-subunit (Fig. 4a). Within this insertion lies the regulatory cysteine pair γCys233–γCys239 (240 and 246 in spinach) that is capable of forming a disulphide bridge. In this way the redox state of the enzyme is communicated to the element of the F_1_ domain that couples conformational changes in the catalytic head to rotary motion^2^. To establish how the *Chlamydomonas* ATP synthase differs from the land-plant enzyme, we compared the insertion and its contacts with the β-subunit in the oxidised and reduced spinach structures and in our consensus model (Fig. 4a-c).

**Fig. 4.**
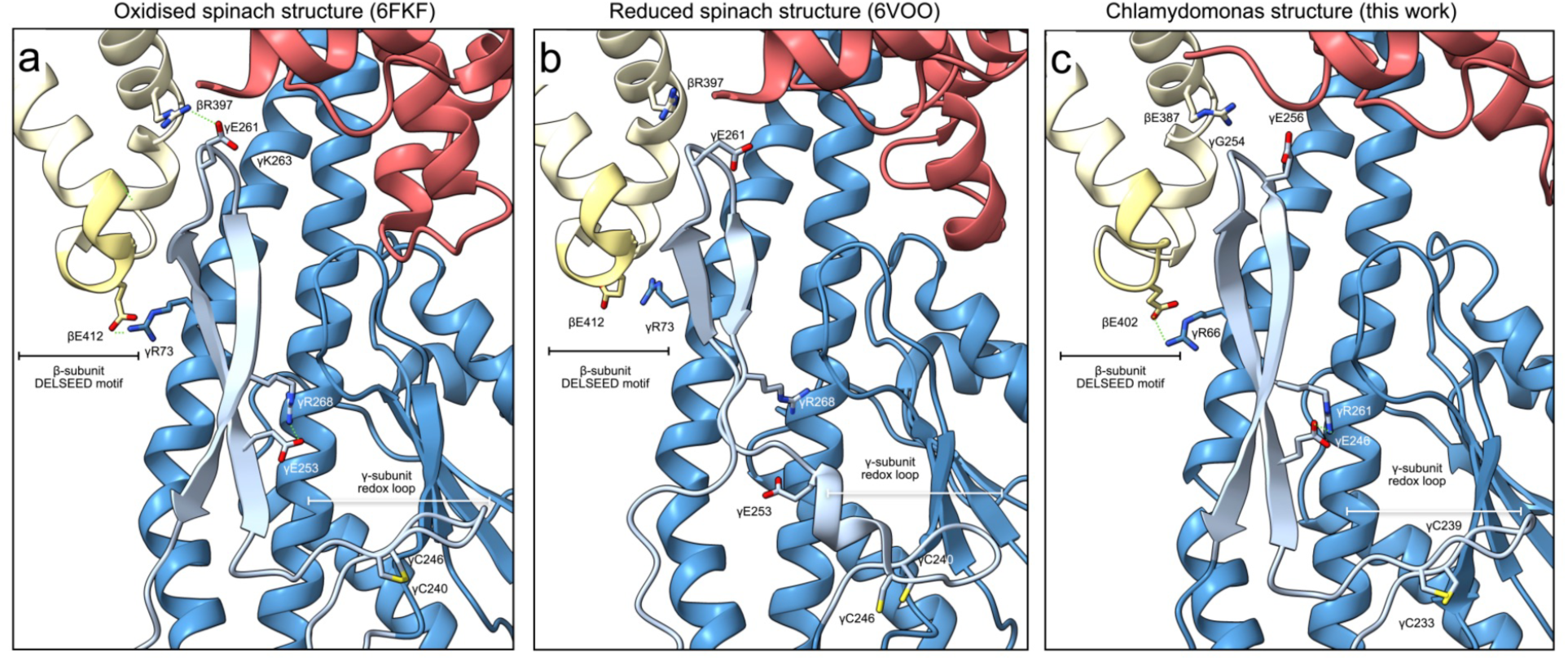
The γ-subunit redox loop retains the disulphide brace but lacks the DELSEED chock in *C. reinhardtii*. Matched views of the γ redox insertion (light blue) and the DELSEED lever of the neighbouring β-subunit (yellow) for ***a***, oxidised spinach (PDB 6FKF^12^, 3.15 Å), ***b***, reduced spinach (PDB 6VOO^11^, 3.05 Å) and ***c***, the *Chlamydomonas* enzyme (this work). The intramolecular brace (γGlu253–γArg268 in spinach; γGlu246–γArg261 in *Chlamydomonas*), the chock (γGlu261–βArg397 in spinach) and the intermolecular tether (γArg73–βGlu412 in spinach; γArg66–βGlu402 in *Chlamydomonas*) are shown as sticks, with the redox-sensitive cysteines (γCys240/γCys246 in spinach; γCys233/γCys239 in *Chlamydomonas*) highlighted. The chock is formed in *a*, released on reduction in *b*, and precluded in *c*, where the position of the chock glutamate is occupied by γGly254 and the conserved partner βArg387 is left unengaged. Interatomic distances are listed in Extended Data Table 3 and buried surface areas in Extended Data Table 4.

In the oxidised spinach enzyme (Fig. 4a; PDB 6FKF)^12^, the γCys240–γCys246 disulphide within the first hairpin of the redox loop is modelled, the two sulphur atoms (Sγ) lying 2.7 Å apart, and the second hairpin is held rigid by an intramolecular brace between γGlu253 and γArg268 (3.7 Å). This brace presents the second hairpin to the DELSEED lever of the adjacent, nucleotide-free (empty) β-subunit through two distinct anchors. The first is a redox-sensitive chock, in which γGlu261 at the tip of the hairpin forms a salt bridge with the conserved βArg397 (3.8 Å); βArg397 lies on the DELSEED-bearing helix immediately upstream of the motif proper (βDELSEED, residues 411 to 417). The second is a tether, in which γArg73 of the N-terminal γ-helix engages βGlu412 within the DELSEED motif (3.7 Å). Reduction of the disulphide bridge abolishes both. In the reduced spinach enzyme (Fig. 4b; PDB 6VOO)^11^ the disulphide in the first hairpin is broken (Sγ–Sγ 4.7 Å), the brace collapses (γGlu253–γArg268, 9.0 Å), and the chock and tether separate to 6.8 and 6.5 Å respectively, releasing the second hairpin from the DELSEED lever (Extended Data Table 3). These structures define the three sets of side-chain contacts through which the redox state of the γCys240–γCys246 pair is transmitted to the catalytic β-subunit.

The *Chlamydomonas* enzyme preserves the general structure and conformation of the oxidised redox loop but not its output (Fig. 4c). The two sulphur atoms (Sγ) of γCys233 and γCys239 lie 3.4 Å apart, consistent with a disulphide bond yet clear density between these residues is absent. However, since disulphides are prone to radiation damage, with their density disappearing in response to a fluence of as little as ∼5 *e*^-^/Å^2^, these atoms most likely represent a disulphide that has broken during imaging^23^. Consistent with this we find the brace observed in the oxidised spinach structure is present, γGlu246 pairing with γArg261 at 3.4 Å, the direct equivalents of spinach γGlu253 and γArg268, and the tether is also fully retained, γArg66 engaging βGlu402 at 2.8 Å. The chock, however, cannot form. The hairpin tip, KEGK in spinach (γLys260–γGlu261–γGly262–γLys263), is replaced by KGGE in *Chlamydomonas* (γLys253–γGly254–γGly255–γGlu256), a substitution that places a glycine, γGly254, at the position occupied by the chock glutamate γGlu261. The partner arginine is conserved and correctly positioned (βArg387, the equivalent of spinach βArg397), and the γGly254 main chain approaches it to within 3.9 Å, yet no side chain is available to complete the interaction (the γGlu256 lies at 5.2 Å distance). The alga therefore assembles the oxidised redox-loop structure without projecting a carboxylate onto the DELSEED lever.

Buried-surface-area analysis quantifies this uncoupling (Extended Data Table 4). The redox insertion buries a comparable total area against the β-subunit in all three structures, between 326 and 370 Å², so the loop occupies the interface irrespective of redox state. The discriminating quantity is the buried side chain of the chock residue: γGlu261 buries 28 Å² of side-chain surface against βArg397 in the oxidised spinach enzyme, this value falls to 17 Å² on reduction, and it is zero in *C. reinhardtii*, where the residue is a glycine. The DELSEED arginine mirrors the trend, its buried side-chain area declining from 54 Å² in the chocked oxidised state to between 37 and 40 Å² when the chock is released or, in the algal enzyme, structurally precluded. At the decisive contact the *C. reinhardtii* interface therefore resembles the reduced spinach interface more than the oxidised one, even though it carries a brace more typical of the oxidised spinach enzyme.

## Discussion

The structure of the *C. reinhardtii* ATP synthase reveals a chloroplast motor that departs from the vascular-plant paradigm at its most fundamental bioenergetic parameter. The previously reported structure modelled a 14-membered ring into a lower-resolution density of the F_o_ domain^16^. Here, with much higher resolution we can definitively show the structure of a 13-membered c-ring, meaning that the *C. reinhardtii* ATP synthase translocates fewer protons per revolution than the c_14_ ring of spinach, and so lowers the H^+^/ATP ratio from 4.67 to 4.33. It is striking that a change of only three interfacial residues is sufficient to alter the c-ring stoichiometry, and it demonstrates that only a small number of mutations is required to shift the H^+^/ATP ratio of a chloroplast ATP synthase; this raises the possibility that stoichiometry has been tuned during the evolution of different photosynthetic lineages in response to their particular energetic requirements. The same principle may offer a route to the targeted engineering of the proton efficiency of photosynthesis^24^.

The Calvin-Benson-Bassham cycle consumes ATP and NADPH in a ratio of 3:2, yet LET supplies these metabolites in a fixed proportion set by the proton budget of the chain^25^. For every two electrons transferred from water to NADP^+^, two protons are deposited in the lumen at photosystem II and four more through the Q-cycle of the cytochrome *b*_6_*f* complex; totalling six protons per NADPH^26^. The ATP recoverable from this budget depends directly on the H⁺/ATP ratio of the synthase, and therefore on c-ring size. A c_14_ ring, with a ratio of 4.67, converts the six protons into 1.28 ATP per NADPH; the c_13_ ring reported here, with a ratio of 4.33, yields 1.39. Neither meets the value of 1.5 demanded by carbon fixation, but the algal stoichiometry halves the ATP deficit that the vascular-plant enzyme would leave, from roughly 0.22 to 0.11 ATP per NADPH. In *C. reinhardtii* this shortfall is thought to be met by a combination of the alternative pathways CET, PCET and CMET, which could generate additional ATP without the net reduction of NADP⁺^14,15^. The carbon-concentrating mechanism raises the ATP demand further still^13^, and the flux through these pathways scales accordingly. Because the saving conferred by the smaller ring arises at the level of ATP yield per unit of LET, it lowers the electron transfer through the alternative pathways required to balance a given demand in the same proportion, to approximately half of the c_14_ value under the baseline stoichiometry of the cycle. The smaller c_13_ ring is thus, in part, a structural adaptation to the higher energetic demands of the carbon-concentrating mechanism. It eases, but does not remove, the alga’s reliance on alternative electron transfer pathways, since the surcharge imposed by the carbon-concentrating mechanism holds the ATP demand above what the change in ring size alone can supply.

Our *C. reinhardtii* structure sheds light not only on the stoichiometry of proton transfer but also on the pathway that carries it. We trace an ordered-water relay linking the two aqueous half-channels of the a-subunit to the protonatable carboxylates at the loading and unloading sites of the c-ring. Because these waters are resolved as discrete density, rather than inferred from the disposition of the surrounding side chains, the proton wire rests on a direct structural footing. The continuous hydrogen-bonded chain provides a route for Grotthuss propagation of the transmembrane proton current, in which charge is relayed from one water molecule to the next rather than carried bodily by a single hydronium ion, which accounts for the anomalously high mobility of protons in water. Directionality is imposed by an a-subunit triad, Arg180, Asn184 and Gln218, that keeps the two half-channels insulated from one another. The ordered-water relay is therefore not an incidental feature of the Fₒ domain but the element that couples transmembrane proton flow to protonation and release at successive c-subunits, and hence to rotation of the ring. A comparable network has recently been described in the phylogenetically distant bacterium *P. aeruginosa*, underlining the deep conservation of Grotthuss proton transfer across the ATP synthase family^21^.

Our data also provide a structural basis for the distinctive regulatory properties of the C. reinhardtii enzyme reported spectroscopically^17^. Unlike the vascular plant chloroplast ATP synthase, oxidation of the γ-subunit thiols in *C. reinhardtii* does not appreciably attenuate activity in the dark. Our structure reveals the basis for this divergence: although the algal enzyme adopts a redox-loop conformation equivalent to that of the oxidised spinach enzyme, a single substitution removes the DELSEED chock^11,12^. The metabolic logic behind the adaptation follows from the ways in which the algal chloroplast differs from that of a vascular plant. A vascular plant is specialised to export photosynthetically assimilated carbon to sustain its non-photosynthetic tissues^27,28^. *C. reinhardtii*, by contrast, is a unicellular alga that can grow not only autotrophically but also heterotrophically and mixotrophically on acetate^29^. Supplied with acetate, the alga uses it both catabolically, to generate ATP through mitochondrial respiration, and anabolically, as a carbon skeleton for the synthesis of lipids, sugars, starch and amino acids in the chloroplast. The chloroplastic arm of this metabolism yields reduced carriers whose electrons enter the thylakoid plastoquinone (PQ) pool through the type II dehydrogenase NDA2 and succinate dehydrogenase^30,31^, and are then passed to O_₂_ by the plastid terminal oxidase (PTOX). This chlororespiratory flux maintains chloroplast redox balance and is essential for growth on acetate^17,32,33^. Because association of PTOX with the thylakoid membrane is pH dependent^34^, the pmf that the algal enzyme is able to generate in the dark, a consequence of the modified γ-subunit, may set the lumenal conditions under which chlororespiration proceeds, and so represent a crucial enabling adaptation to the mixotrophic lifestyle^17,32^.

Taken together, our structure resolves unique features that establish a structural framework for the bioenergetics of the green algal chloroplast and moreover invite a re-examination of how the balance between electron transfer and carbon fixation is set in an organism that concentrates its own CO_2_. The latter should be considered when transplanting algal-type carbon concentrating mechanisms into vascular plants^13^.

## Methods

### Thylakoid isolation and protein purification

Thylakoid and protein preparation were performed as described previously^35^ with the following modifications. *C. reinhardtii* UVM4 was cultured in 2 L TAP medium under a 14 h/10 h day–night cycle at 60 µmol photons m^-2^ s^-1^ white LED light at 20 °C for 3 days, and cells were collected by centrifugation at 4,000 × g for 15 min. The pellet was resuspended in 10 mM HEPES pH 7.5, 0.33 M sucrose, 10 mM EDTA, 10 mM NaF, 1.5 mM KCl, supplemented with EDTA-free protease inhibitor (Merck) and DNase, and lysed by two passes through a French press at 8,000 psi. Lysate was clarified at 3,000 × g for 10 min at 4 °C and thylakoids collected at 235,418 × g, for 50 min at 4 °C. Thylakoids were resuspended to 2 mg mL^-1^ chlorophyll in 10 mM HEPES pH 7.5, 10 mM EDTA, 1.5 mM KCl, solubilised with 2% (w/v) α-DDM for 30 min at 4 °C in the dark, and insoluble material removed at 185,976 × g for 45 min. Solubilised complexes (1.5 mL) were loaded onto 10 mM HEPES pH 7.5, 0.85 M sucrose, 0.009% α-DDM density gradients and centrifuged for 20 h at 124,513 × g at 4 °C.

### Cryo-EM sample preparation and data acquisition

Purified ATP synthase was concentrated to ∼5 mg mL^-1^ (100 kDa MWCO) and exchanged into 20 mM HEPES pH 7.5, 10 mM NaCl, 1 mM EDTA, 0.01% α-DDM. Grids (Quantifoil R2/1 Cu 200 mesh) were glow-discharged for 6 s at 15 mA; 3 µL of sample was applied and vitrified in a Vitrobot Mark IV (95% humidity, 277 K, 3 s blot, blot force 3). Data were collected at the National Synchrotron Radiation Centre SOLARIS (Krakow) on a Titan Krios G3i at 300 kV, ×105,000 magnification and 0.84 Å per pixel, using a K3 detector with a BioQuantum energy filter (20 eV slit) in counting mode. In total 17,815 movies (40 frames; total dose 41.5 e^-^ Å^-2^) were recorded over a defocus range of -1.0 to -2.0 µm.

### Cryo-EM data processing and model building

Processing was performed in cryoSPARC v4.7.0^36^. Movies were patch motion-corrected and CTF-estimated. Particles were picked with a Topaz model trained on an initial clean set, extracted (512-pixel box) and cleaned by two-dimensional classification and ab-initio reconstruction. Non-uniform refinement with per-particle CTF and defocus optimisation, Ewald-sphere correction, and reference-based motion correction yielded a 2.2 Å consensus map from 235,203 particles. A high-resolution composite map was assembled in ChimeraX ^37^ from local refinements of the c-ring, F_1_, γ+ε and δ+a+bb’ regions. Rotary states were separated by three-dimensional classification (four classes, 7 Å filter), giving state 1, state 2 and substates 3.1 and 3.2; per-state F_1_-masked local refinements reached 2.3–2.7 Å, and an F_o_-masked refinement of the combined state-3 particles reached 2.7 Å with defined waters. The particles belonging to state 3 were further classified in 3D (four classes, 6 Å filter) with a focus mask over the F_1_ portion to reveal two conformations of the α subunit.

The consensus model was built from an AlphaFold3^38^ prediction in ChimeraX and Coot 2.9.8.96^39^. The F_o_ domain had speckles of density on its surface that corresponded to lipids binding to its hydrophobic surface. As these can look like waters, waters were only built when they were clearly bound in the interior of the domain. Once model building was complete it was refined in PHENIX^40^ real-space refinement and validated with MolProbity^41^; models for related maps were derived from the consensus model.

### Structural analysis

Ring geometry of the *C. reinhardtii* c_13_ and spinach c_14_ (PDB 6FKF) structures was analysed with gemmi and NumPy. Subunit centres of mass were computed from Cα coordinates and the ring plane defined by singular-value decomposition; inner and outer wall radii were taken as the minimum and maximum in-plane Cα distance from the ring axis, and angular and neighbour spacings computed in the fitted plane. Helix tilts were obtained by principal-component analysis of the inner and outer transmembrane helices. Interface buried surface area was calculated with FreeSASA (probe radius 1.4 Å) over all protomer pairs. Inter-state rotations were measured after superposition of the α_3_β_3_ head, and γ redox-loop geometry compared with the oxidised (PDB 6FKF) and reduced (PDB 6VOO) spinach models. Interatomic distances at the redox interface were measured between polar side-chain atoms with gemmi, and buried surface areas were computed with FreeSASA using the Lee–Richards algorithm and a 1.4 Å probe, as the difference in solvent-accessible surface between the isolated γ- and β-subunits and their complex; the contacting β-subunit was that making the closest approach to the γ redox insertion in each structure (β-subunit B in 6FKF, E in 6VOO and U in the consensus model). Sequence alignment used BLOSUM62 in Biopython, and superpositions and molecular graphics were prepared in PyMOL v2.5.

## Acknowledgements

M.P.J. acknowledges funding from the Leverhulme Trust (RPG-2021-345) and the BBSRC (UKRI1945). C.N.H. and K.L. acknowledge European Research Council Synergy Award 854126. A.H. is funded by a Royal Society University Research Fellowship (URF\R1\191548 & URF\R\241006). J.N.B. is a UKRI Future Leader Fellow (MR/T040742/1 & MR/Z000084/1). S.P. acknowledges support from the Faculty of Biochemistry, Biophysics and Biotechnology under the Strategic Programme Excellence Initiative at Jagiellonian University (WBBiB.2.2.2025.11). We acknowledge the University of Sheffield Cryo-Electron Microscopy Facility, PLGrid (ACK Cyfronet AGH) for computational resources (PLG/2025/018653) and the Polish Ministry of Science and Higher Education project 1/SOL/2021/2 supporting the National Synchrotron Radiation Centre SOLARIS.

## Author contributions

K.L. and S.P. performed cryo-EM grid screening, data processing and model building; K.H.R., H.T. and M.S.P. contributed to sample preparation and structural analysis; M.R. performed cryo-EM data collection; C.N.H., A.H., J.N.B. and M.P.J. supervised the work; M.P.J., K.L., A.H., and J.N.B. wrote the manuscript with input from all authors.

## Data availability

The atomic model and cryo-EM map of the *C. reinhardtii* ATP synthase have been deposited in the PDB and EMDB under accession codes PDB 33RO and EMD-59509 (consensus model and the corresponding composite map). The consensus and focused local refinement maps used to generate the composite map were deposited in the EMDB under accession numbers: EMD-59504 (consensus), EMD-59505 (F1 focus), EMD-59506 (c-ring focus), EMD-59507 (sub. γ and ε focus) and EMD-59508 (sub. δ, a, b and b’ focus). Local-refinement and rotary-state maps and models are deposited under codes EMD-XXXX to EMD-XXXX, and PDB IDs XXX to XXX.

---

**Extended Data Fig. 1.**
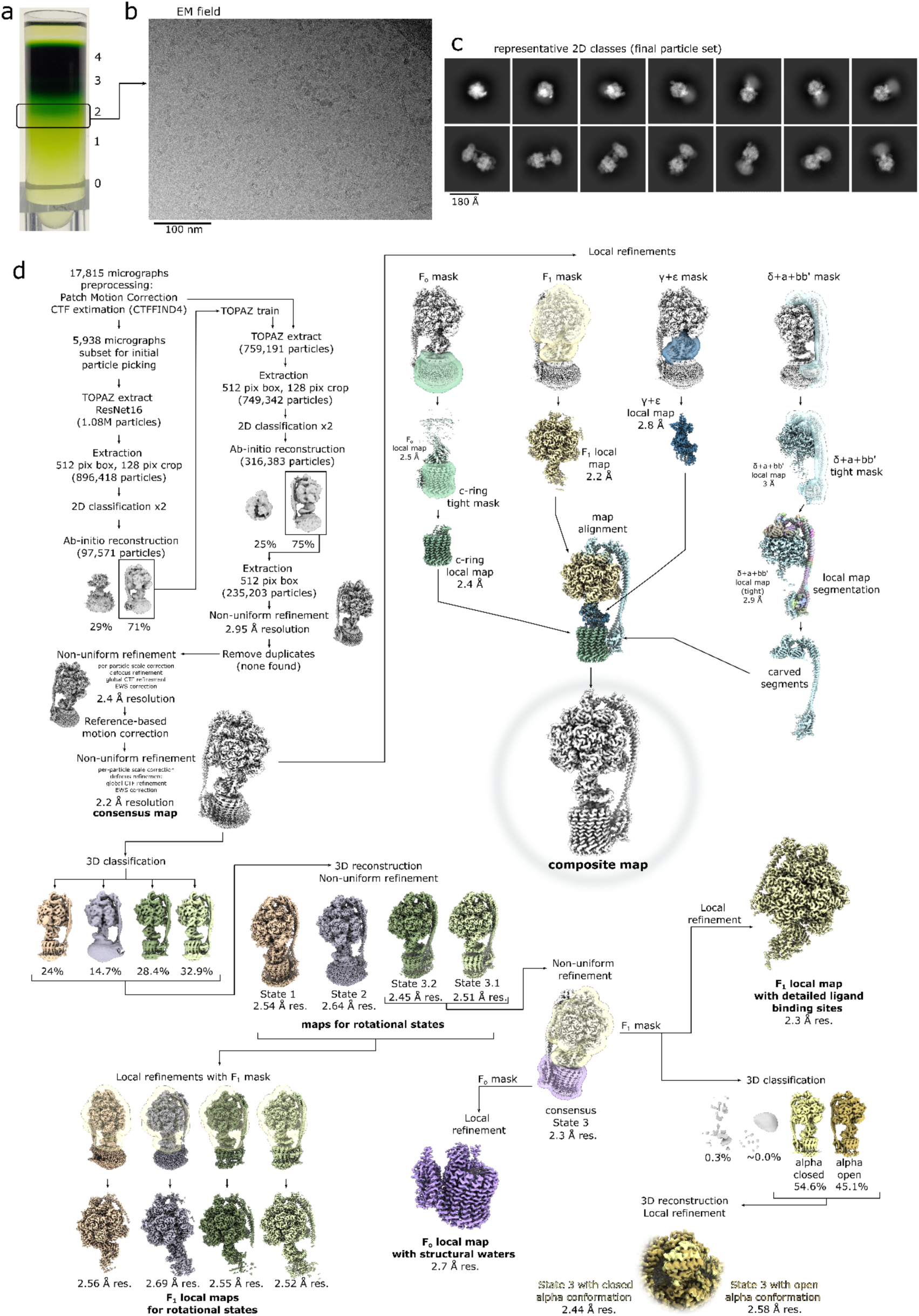
CryoEM data processing workflow: **a,** sucrose gradient ultracentrifugation of α-DDM solubilised thylakoids (see methods), band 2 was selected for cryo-EM without further purification, **b**, representative cryo-EM field; **c,** selected representative 2D class averages for the final particle set; **d,** cryoSPARC processing and the composite map preparation pipeline.

**Extended Data Fig. 2.**
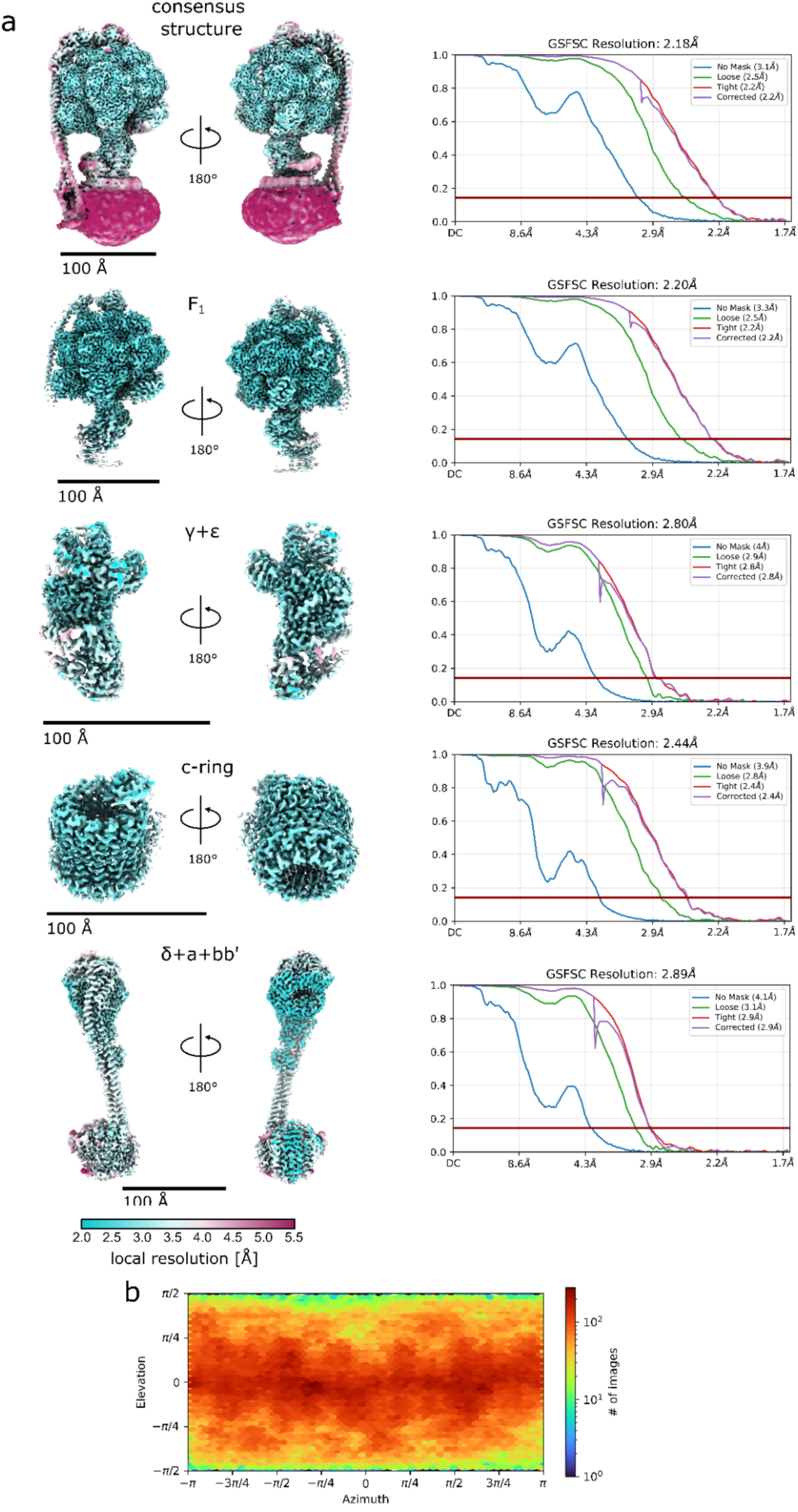
Quality evaluation for the consensus structure cryo-EM map and local refinement maps used to reconstruct the composite map: **a,** left: local-filtered maps coloured by resolution range (scale at the bottom); right: respective Fourier shell correlation curves, dark red lines in the plots indicate FSC = 0.143; **b,** Particle angular distribution heatmap for the consensus structure calculated in cryoSPARC.

**Extended Data Fig. 3.**
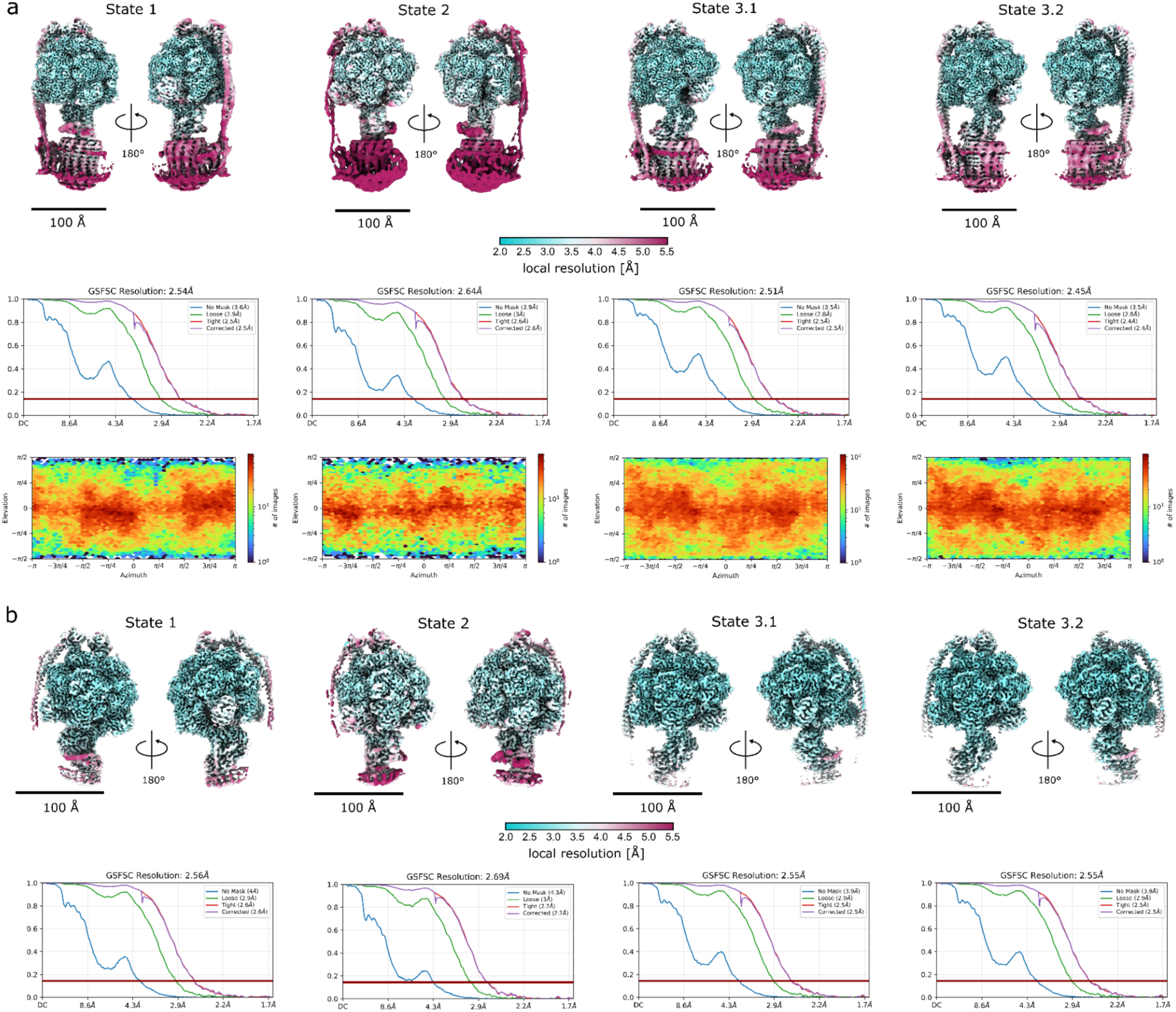
Quality evaluation for the rotational states and their F1-focused maps. **a,** top: local-filtered maps for rotational states coloured by resolution range (scale at the bottom); middle: respective Fourier shell correlation curves, dark red lines in the plots indicate FSC = 0.143; bottom: respective particle angular distribution heatmaps calculated in cryoSPARC **b,** top: local-filtered, F1-focused local-refined maps for rotational states coloured by resolution range (scale at the bottom); bottom: respective Fourier shell correlation curves, dark red lines in the plots indicate FSC = 0.143.

**Extended Data Fig. 4.**
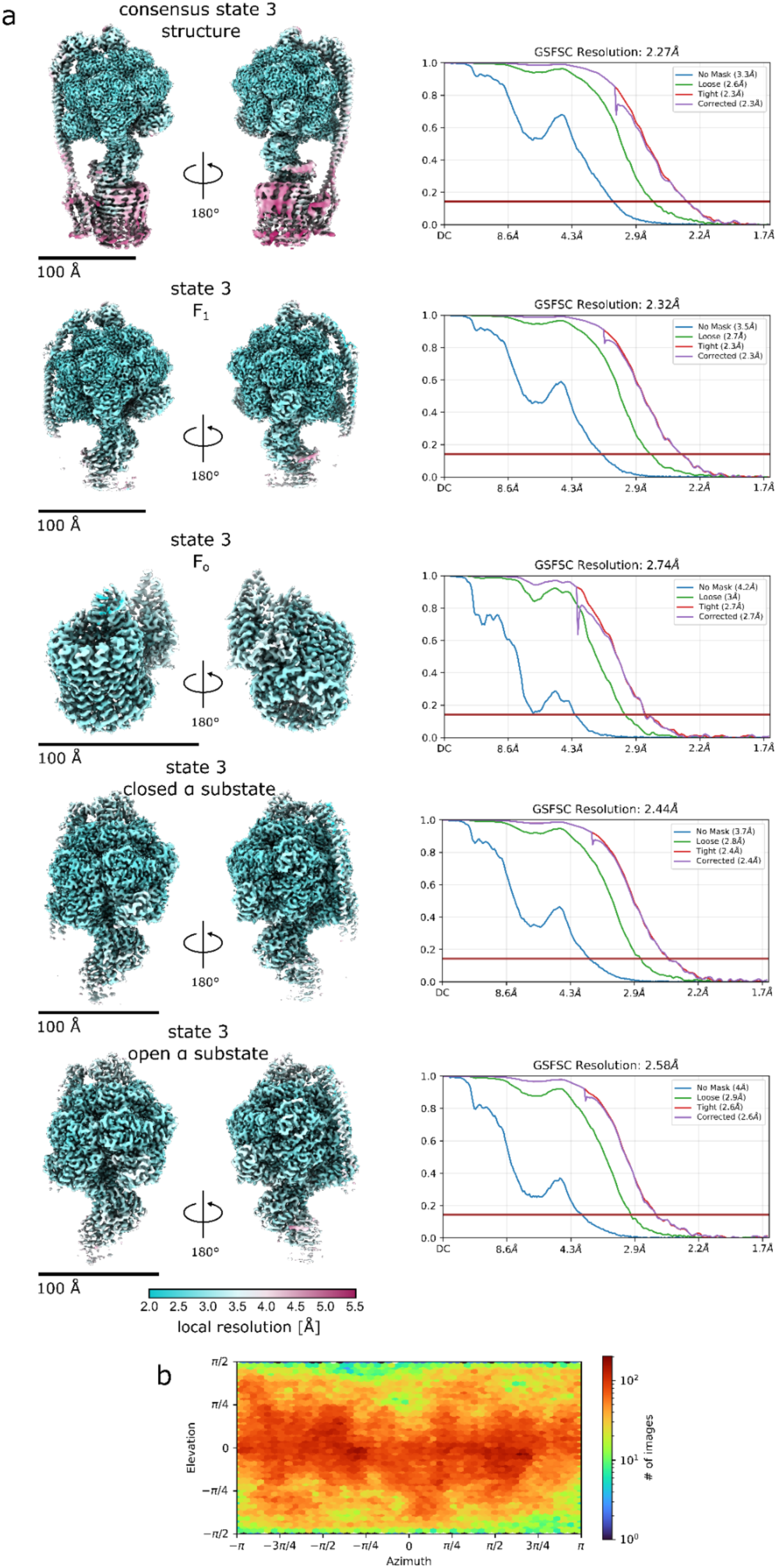
Quality evaluation for the consensus rotational state 3 and focused-refined maps of state 3. **a,** left: local-filtered maps for consensus state 3 map and maps from local refinements (F1, Fo and α-subunit at closed and open conformation) coloured by resolution range (scale at the bottom); right: respective Fourier shell correlation curves, dark red lines in the plots indicate FSC = 0.143; **b,** particle angular distribution heatmap for consensus state 3 map calculated in cryoSPARC.

**Extended Data Fig. 5.**
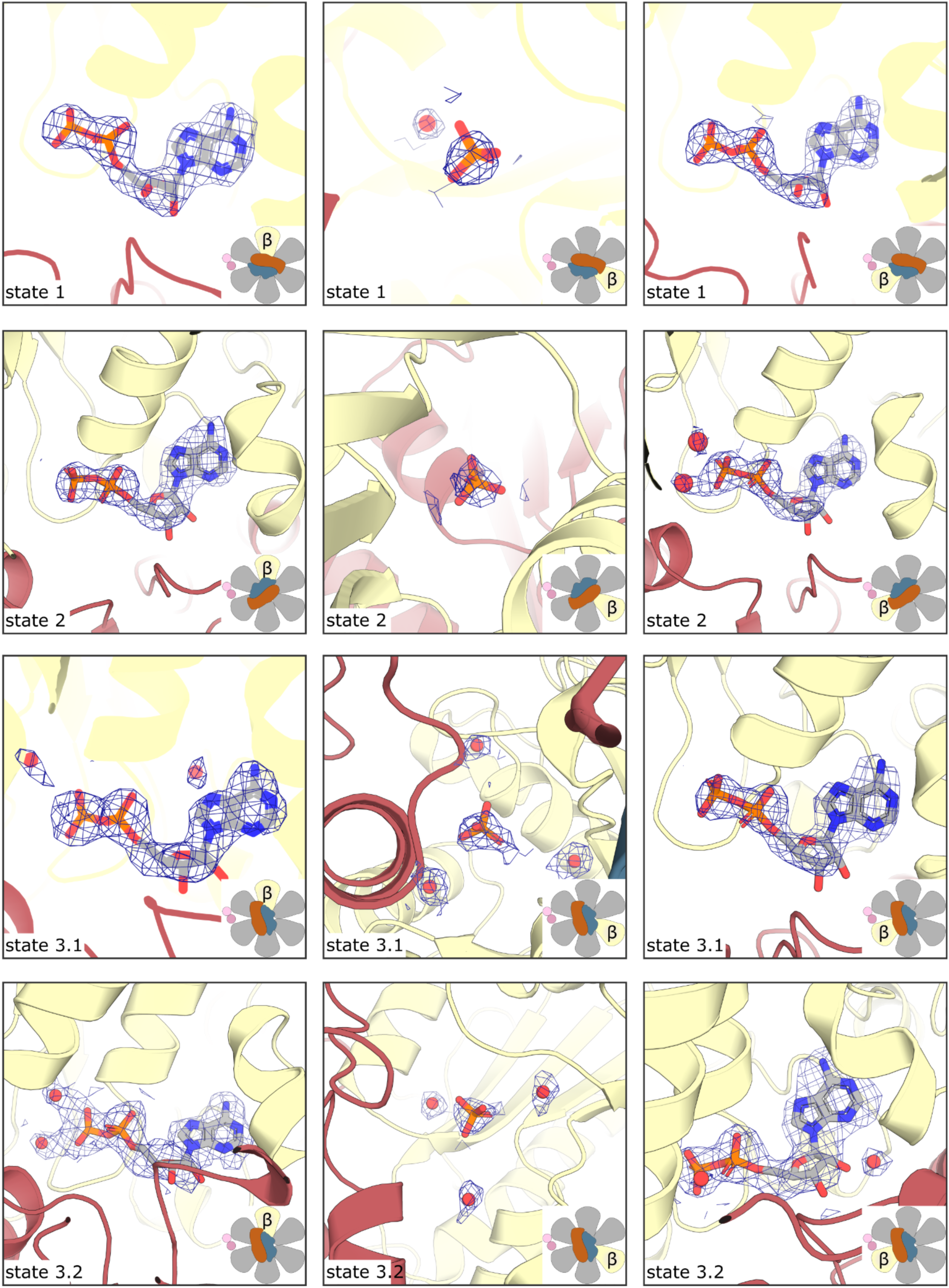
Full comparison of β-subunit binding site occupancies /nucleotide densities in the various substates presented in Figure 1. Waters are shown as red spheres. In the bottom right hand corner of each image the position of the β-subunit illustrated is shown relative to the angle/position of the gamma (blue), epsilon (orange) and stator (b/ b’) subunits (pink circles).

**Extended Data Fig. 6.**
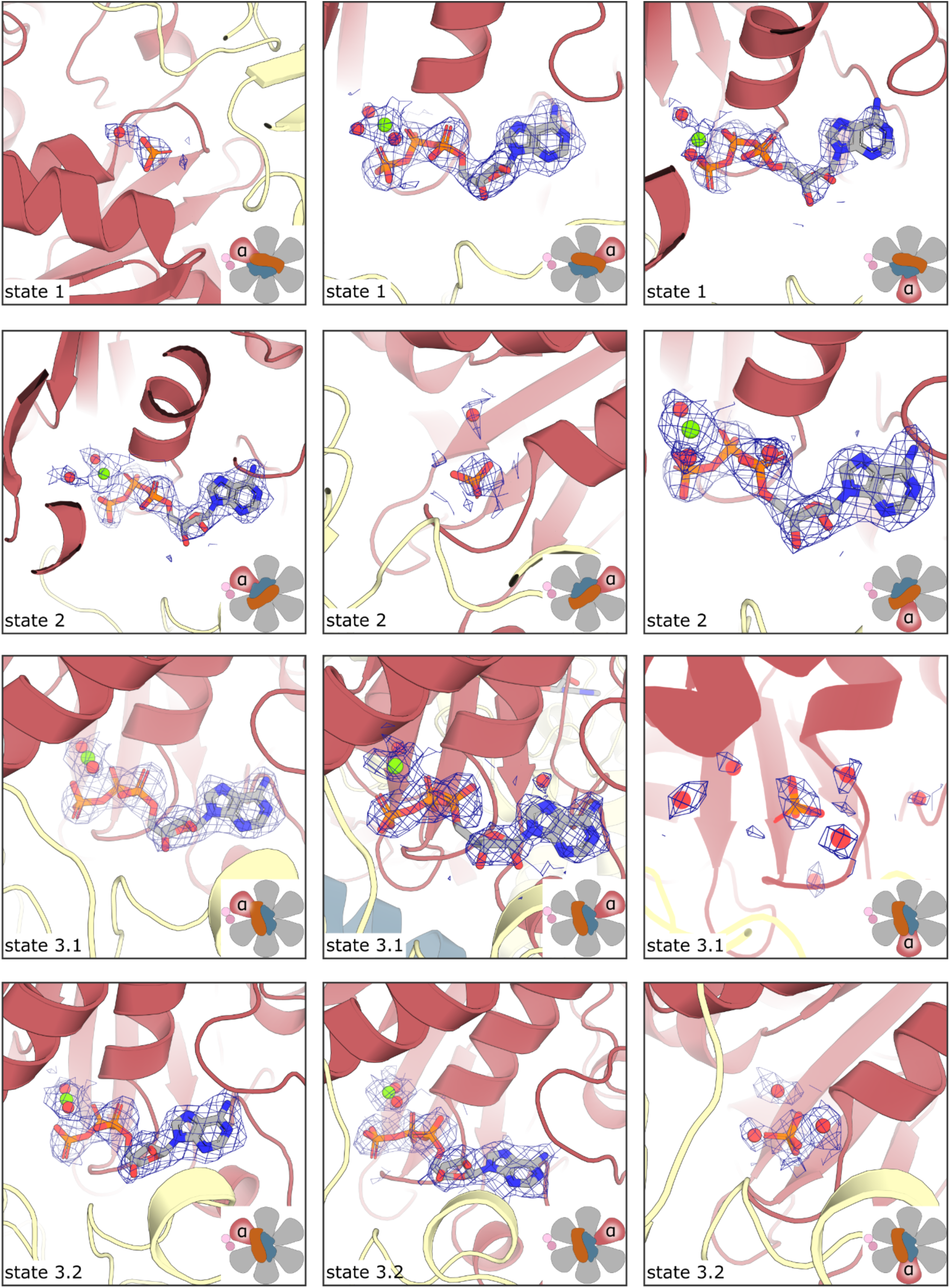
Full comparison of α-subunit binding site occupancies /nucleotide densities in the various substates presented in Figure 1. Waters are shown as red spheres and Mg^2+^ cations as green spheres. In the bottom right hand corner of each image the position of the α-subunit illustrated is shown relative to the angle/position of the gamma (blue), epsilon (orange) and stator (b/ b’) subunits (pink circles).

**Extended Data Table 1.**
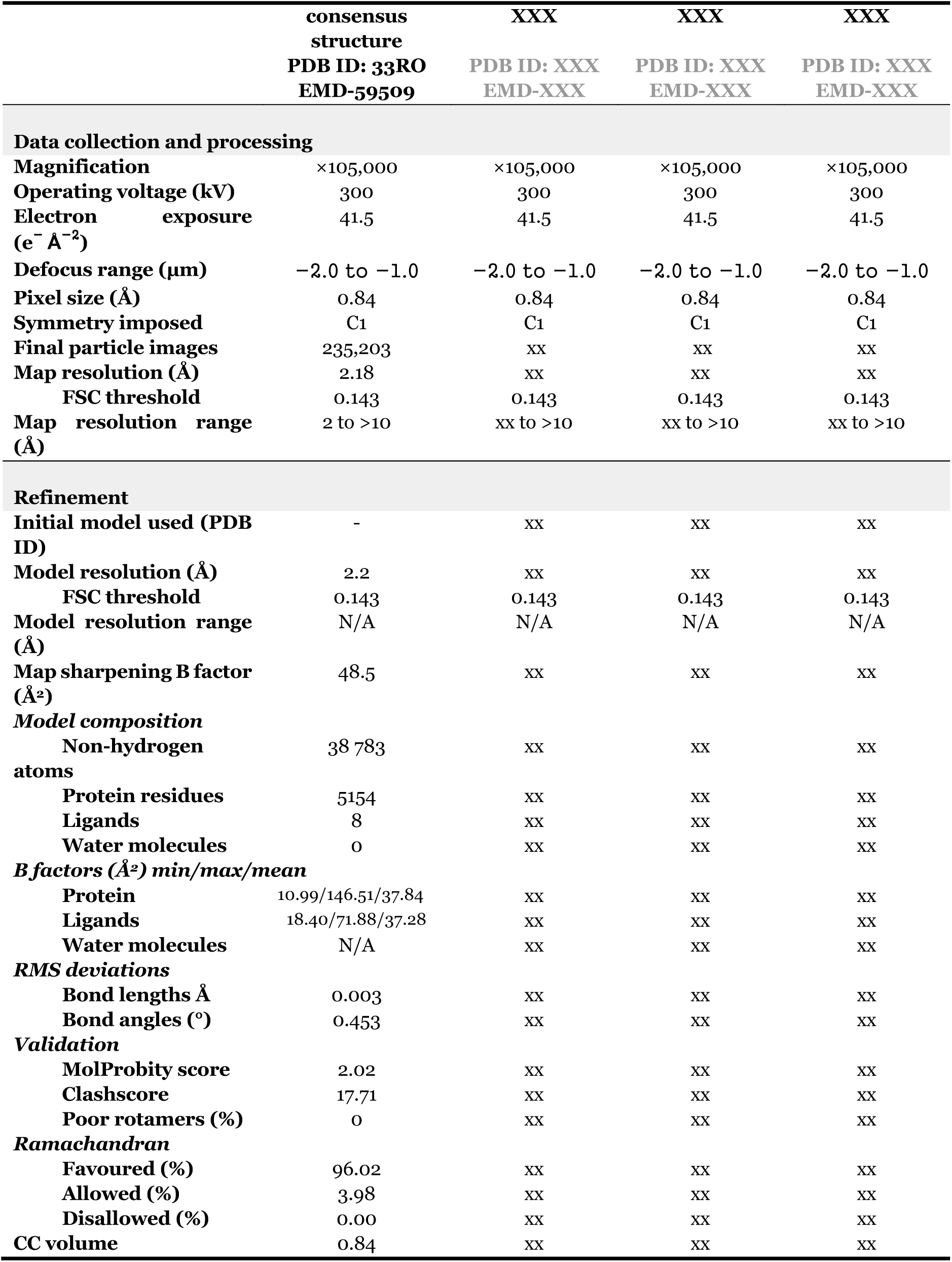
Cryo-EM data collection, refinement and validation statistics.

|  | consensus<br>structure<br>PDB ID: 33RO<br>EMD-59509 | XXX<br>PDB ID: XXX<br>EMD-XXX | XXX<br>PDB ID: XXX<br>EMD-XXX | XXX<br>PDB ID: XXX<br>EMD-XXX |
| --- | --- | --- | --- | --- |
| **Data collection and processing** |  |  |  |  |
| Magnification | ×105,000 | ×105,000 | ×105,000 | ×105,000 |
| Operating voltage (kV) | 300 | 300 | 300 | 300 |
| Electron exposure<br>(e<sup>-</sup> Å<sup>-2</sup>) | 41.5 | 41.5 | 41.5 | 41.5 |
| Defocus range (μm) | -2.0 to -1.0 | -2.0 to -1.0 | -2.0 to -1.0 | -2.0 to -1.0 |
| Pixel size (Å) | 0.84 | 0.84 | 0.84 | 0.84 |
| Symmetry imposed | C1 | C1 | C1 | C1 |
| Final particle images | 235,203 | XX | XX | XX |
| Map resolution (Å) | 2.18 | XX | XX | XX |
| FSC threshold | 0.143 | 0.143 | 0.143 | 0.143 |
| Map resolution range<br>(Å) | 2 to >10 | xx to >10 | xx to >10 | xx to >10 |
| **Refinement** |  |  |  |  |
| Initial model used (PDB ID) | - | XX | XX | XX |
| Model resolution (Å) | 2.2 | XX | XX | XX |
| FSC threshold | 0.143 | 0.143 | 0.143 | 0.143 |
| Model resolution range<br>(Å) | N/A | N/A | N/A | N/A |
| Map sharpening B factor<br>(Å<sup>2</sup>) | 48.5 | XX | XX | XX |
| **Model composition** |  |  |  |  |
| Non-hydrogen<br>atoms | 38 783 | XX | XX | XX |
| Protein residues | 5154 | XX | XX | XX |
| Ligands | 8 | XX | XX | XX |
| Water molecules | 0 | XX | XX | XX |
| **B factors (Å<sup>2</sup>) min/max/mean** |  |  |  |  |
| Protein | 10.99/146.51/37.84 | XX | XX | XX |
| Ligands | 18.40/71.88/37.28 | XX | XX | XX |
| Water molecules | N/A | XX | XX | XX |
| **RMS deviations** |  |  |  |  |
| Bond lengths Å | 0.003 | XX | XX | XX |
| Bond angles (°) | 0.453 | XX | XX | XX |
| **Validation** |  |  |  |  |
| MolProbity score | 2.02 | XX | XX | XX |
| Clashscore | 17.71 | XX | XX | XX |
| Poor rotamers (%) | 0 | XX | XX | XX |
| **Ramachandran** |  |  |  |  |
| Favoured (%) | 96.02 | XX | XX | XX |
| Allowed (%) | 3.98 | XX | XX | XX |
| Disallowed (%) | 0.00 | XX | XX | XX |
| CC volume | 0.84 | XX | XX | XX |

**Extended Data Table 2.** Ring geometry, buried surface area calculations and helix-tilt analysis of Fo rotor.

| | *Chlamydomonas*<br>C13 | Spinach<br>C14 | $\Delta$ |
| --- | --- | --- | --- |
| **Ring geometry** |  |  |  |
| Subunit COM radius | 21.19 Å | 21.63 Å | −0.44 Å |
| Inner wall radius (min Cα–axis) | 12.78 Å | 13.08 Å | −0.30 Å |
| Outer wall radius (max Cα–axis) | 30.65 Å | 31.24 Å | −0.59 Å |
| Mean neighbour COM–COM distance | 10.14 Å | 9.63 Å | +0.52 Å |
| Angular subunit spacing | 27.69° | 25.71° | +1.98° |
| **Helix-tilt analysis** |  |  |  |
| Inner helix tilt | 5.20 ± 0.14° | 5.16 ± 0.14° | +0.04° |
| Outer helix tilt | 1.19 ± 0.27° | 0.53 ± 0.32° | +0.67° |
| Inner helix length | 45.3 Å | 41.1 Å | +4.2 Å |
| Outer helix length | 44.1 Å | 43.5 Å | +0.6 Å |
| **Buried surface area** |  |  |  |
| Total BSA per interface | 3498 ± ~50 Å<sup>2</sup> | 3608 ± 51 Å<sup>2</sup> | −110 Å<sup>2</sup> |
| BSA per subunit | 1749 Å<sup>2</sup> | 1804 Å<sup>2</sup> | −55 Å<sup>2</sup> |

**Extended Data Table 3.** Key interface distances (Å).

| Contact (spinach numbering) | Oxidised spinach 6FKF | Reduced spinach 6VOO | *Chlamydomonas* (this work) |
| --- | --- | --- | --- |
| γCys240–γCys246 disulphide (Sγ–Sγ) | 2.7 | 4.7 | 3.4 |
| Brace γGlu253–γArg268 | 3.7 | 9.0 | 3.4 |
| Chock γGlu261–βArg397 | 3.8 | 6.8 | absent (γGly254) |
| Tether γArg73–βGlu412 | 3.7 | 6.5 | 2.8 |

*Chlamydomonas* equivalents: γCys233/ γCys239, γGlu246– γArg261, γGly254 / βArg387, γArg66– βGlu402.

**Extended Data Table 4.** Buried surface areas at the redox loop–DELSEED interface (Å²).

| Structure | Redox insertion (hp1+hp2) | Chock-site side chain (γGlu261 / γGly254) | DELSEED arginine side chain (βArg397 / βArg387) |
| --- | --- | --- | --- |
| Oxidised spinach (6FKF) | 370 | 28 | 54 |
| Reduced spinach (6VOO) | 326 | 17 | 37 |
| *Chlamydomonas* (this work) | 343 | 0 | 40 |

Buried area is the loss of solvent-accessible surface on complex formation, computed per side. The redox insertion spans $\gamma$ 236–270 (spinach) and $\gamma$ 229–263 (*Chlamydomonas*). The chock-site value is the side-chain area buried by the $\gamma$ residue at motif position 2, a glutamate in spinach and a glycine in *Chlamydomonas*; the glycine value is zero by definition.

